# Archival data storage using DNA proximity graphs

**DOI:** 10.64898/2026.10.12.758707

**Authors:** Eyal Perry, Jan O. Tiepelt, Joseph M. Jacobson

**Affiliations:** Massachusetts Institute of Technology, Cambridge, MA 02139, USA

## Abstract

DNA offers a promising medium for long-term archival data storage, but writing information into DNA remains costly. Here we introduce SPAN (Synthesis-free Proximity-graph Archiving in Nucleic acids), a synthesis-independent storage architecture in which information is written as a physical pattern and encoded in the spatial relationships of randomly barcoded DNA molecules. Barcodes are deposited onto an optically patterned surface, and an enzymatic reaction records neighboring barcode pairs. Sequencing these pairs produces a proximity graph from which the information is decoded. We demonstrate this principle by writing a 136-bit pattern using photolithography and recovering 98.5% of the bits. Our cost analysis estimates 10X lower writing costs than any existing DNA storage method. Using established commercial optical writing technologies, we project a further five-order-of-magnitude reduction to below flash memory costs. Although readout is more expensive, this trade-off is attractive for deep archival storage, where information is written once and rarely, if ever, retrieved.

## Introduction

The global datasphere is projected to exceed 380 zettabytes (1 ZB = 10^12^ GB) by 2028 (*1*). Increasingly, data are generated without human involvement: Internet of Things (IoT) devices produce data faster than people, and the recent proliferation of autonomous artificial intelligence (AI) agents could accelerate this trend further. Data production has already outpaced the growth of magnetic, optical and solid-state storage media, reducing the fraction of it that can be retained.

It is estimated that 80% of global data storage is archival, accessed rarely or never (*2, 3*). Read probability declines with age, becoming negligible after a decade (Write Once, Read Never; WORN). Yet scientific, legal and cultural records may require retention for decades, lifetimes or indefinitely (*4*). Conventional media last 10-25 years, requiring repeated migration and replacement, with recurring hardware, power, and maintenance costs. Maintaining an exabyte-scale (10^9^ GB) data center costs an estimated $1 billion per decade (*5*). This presents an opportunity for a technology that is inexpensive to write and store for a lifetime, even if read costs are high.

Molecular storage is a promising solution for deep archival storage, potentially storing exabytes of information at a fraction of the footprint, power and cost of conventional media (*6 –8*). DNA, the information-storage medium of biology, is the most developed molecular storage framework (*9 –13*). Its theoretical storage density is 4.5 × 10^11^ GB per gram (*14*), approximately six orders of magnitude denser than the densest electronic media (*15*), and up to 4.3 × 10^10^ GB/g has been achieved experimentally (*16, 17*). At such densities, data currently requiring hangar-sized facilities could fit on a standard table. DNA is also chemically stable for millennia (*18*), as demonstrated by the recovery of million-year-old mammoth DNA (*19*), and requires no power at rest or periodic migration.

The main obstacle to adoption of DNA storage is the writing cost. Most approaches rely on data-specific DNA synthesis, in which information is encoded through the iterative addition of nucleotides, either by conventional phosphoramidite chemistry or enzymatic synthesis (*20*). The most promising strategy for reducing synthesis costs has been miniaturization and parallelization using semiconductor (CMOS) arrays (*21, 22*), with the most economical reported approach combining enzymatic synthesis with CMOS arrays to reach $31 per megabyte (*23*). This strategy has an upper limit: smaller feature size increases chemical crosstalk and diffusion, which degrade synthesis fidelity (*24*). More recently, synthesis-free writing approaches have emerged. Inspired by movable-type printing, these methods use prefabricated DNA building blocks that are synthesized independently of the data being stored. These blocks are then either combinatorially assembled by enzymatic ligation (*25*), or used to direct enzymatic methylation on a universal template (*26*). Although these approaches eliminate the need for data-specific DNA synthesis, writing costs remain higher than CMOS, and further reductions depend on improvements in reaction chemistry and liquid-handling throughput.

Optical writing is a well-established approach to data storage. There, a focused laser forms pits, voids or nanostructures, or alters a material’s refractive index. It ranges from mass-produced optical discs (CD, DVD, Blu-ray), which degrade over decades, to laser writing in glass and ceramic projected to survive millennia (*27, 28*). Compared with molecular storage, optical writing is fast and inexpensive because its rate is governed by laser modulation and scanning rather than sequential synthesis cycles. Its feature size, however, is constrained by wavelength and material response, with even mature, high-density optical storage technologies such as Blu-ray operating at feature sizes on the order of 100nm, far larger than the ∼2 nm diameter of double-stranded DNA. Optical writing is six orders of magnitude cheaper per bit than DNA synthesis, while storing each bit in nine orders of magnitude more volume (Table S2).

Here we combine the two. We introduce SPAN (**S**ynthesis-free **P**roximity-graph **A**rchiving in **N**ucleic acids), which decouples data writing from DNA manipulation. Unlike conventional DNA storage, which encodes bits in specific sequences, SPAN uses a randomly barcoded library synthesized at a fixed cost, independent of the data written. A bit pattern is optically written onto a planar surface, and the library is adsorbed onto the background but not the written features, so that the data now resides in which barcode landed where. An enzymatic reaction generates products containing barcodes of neighboring molecules. These double-barcode pairs constitute the molecular storage medium. Pairwise proximity information can be used to infer spatial organization (*29, 30*). Proximity graphs of DNA barcodes have been used to recover the spatial organization of tissues (*31*), proteins on single-cell membranes (*32*), whole organisms (*33*) and synthetic bead arrays (*34*). Here we apply this principle to a laser-written bit pattern. We demonstrate SPAN on lithographically patterned chips and evaluate experimental and projected costs. Even at proof-of-concept scale, estimated writing costs are an order-of-magnitude lower than the most economical synthesis-based approach. Established optical writing technologies could reduce the write cost by five more orders of magnitude. This shifts most of the cost from writing to substantially more expensive reading. For data that is written once and read with low probability, and whose reading, if it happens at all, happens decades later, hence on a sequencing cost curve that has fallen exponentially over the last two decades (*35*), this is a favorable trade.

## Results

### Encoding bits as holes in proximity graphs

Information is encoded as a 2D binary pattern on a lattice, where written features represent 1 and bare sites represent 0 (Fig. 1A). Sites are spaced to maintain a connected unwritten background, a requirement for later graph reconstruction. The pattern can be produced using established planar writing technologies, including photolithography, electron-beam lithography or optical-disc mastering, and nothing about this step involves DNA. The essential requirement is selective DNA adsorption onto the unwritten background rather than the written features.

**Fig. 1.**
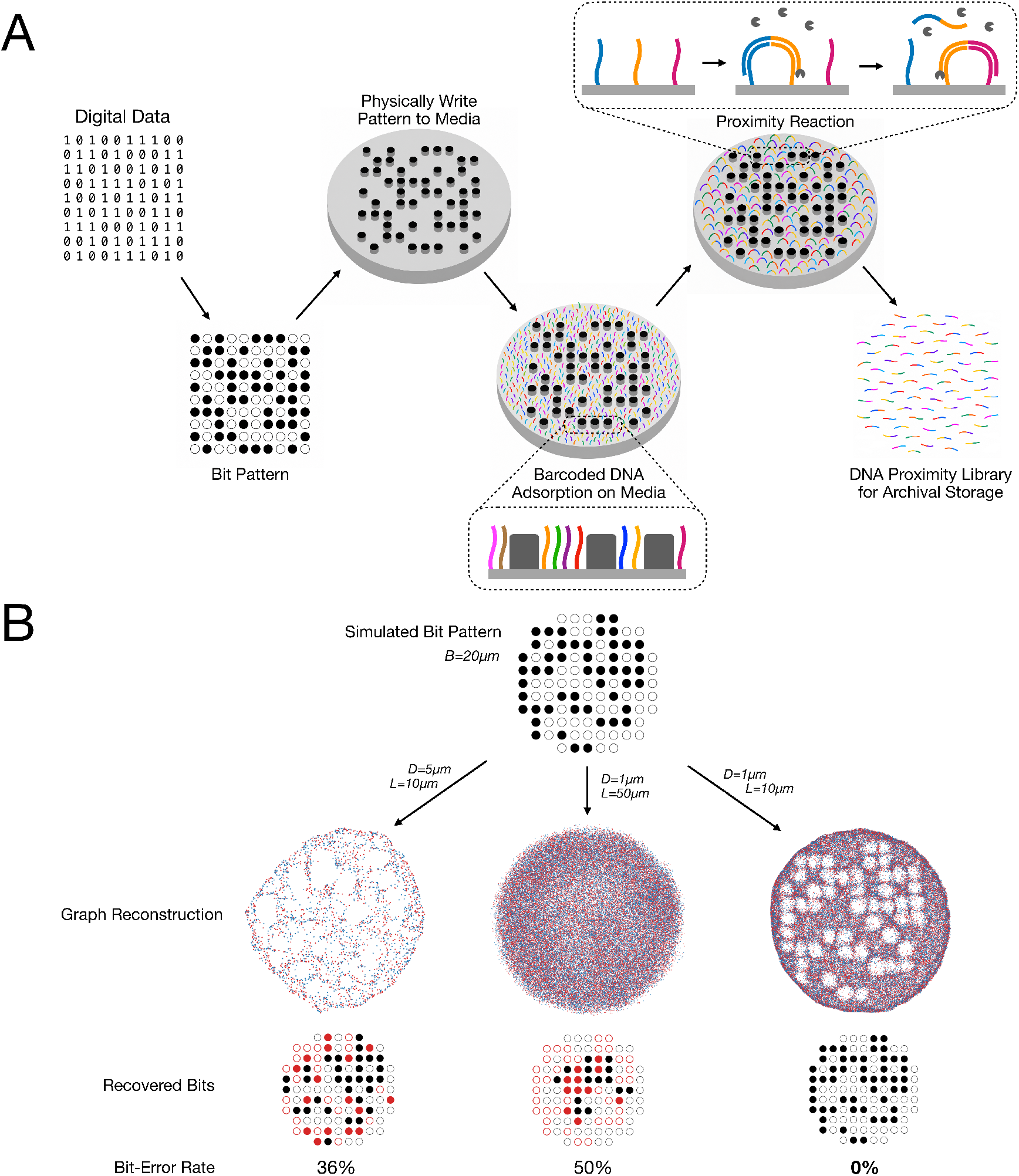
SPAN Principle. (**A**) Digital data is converted into a spatial bit pattern, which is physically written onto a 2D surface, here as bumps at the positions of the 1s. The background around the bumps remains connected, a requirement for later reconstruction. Barcoded DNA molecules are then adsorbed over the surface, binding to the background only and not to the bumps. An enzymatic reaction generates new DNA strands that contain pairs of barcodes that are adjacent on the surface. These barcode-pair strands are harvested from the surface and are the SPAN storage medium. (**B**) A random 101-bit pattern (bit width B=20µm) is simulated. Bipartite proximity graphs are generated at varying node spacings (D=1 or 5 µm) and interaction lengths (L= 10 or 50µm). Red and blue dots represent the two types of nodes in the graph. Each graph is reconstructed and used for bit decoding. Recovered bits are illustrated as black filled circles for true 1s, empty black circles for true 0s, red filled circles for falsely decoded 1s, and empty red circles for falsely decoded 0s. Reconstruction is invariant to rotation and reflection, such that resulting images are manually aligned for visualization. Low molecular density or long interaction range leads to decoding errors, whereas sufficient density and short interactions enable accurate recovery of all 101 bits (0% error rate).

The patterned surface is exposed to a barcoded DNA library. Synthesized once, the same library can serve any data, so its synthesis cost is fixed. Barcode space must be sufficiently complex for individual molecules to be uniquely identifiable: a 25-nucleotide barcode provides 4^25^ (over a quadrillion) possible sequences. Selective adsorption onto the unwritten background creates “holes” in an otherwise uniform molecular layer. The layer must be sufficiently uniform to avoid false bits from unintended voids, while its density must be sufficiently high, as the mean spacing between molecules determines the smallest features that can be resolved.

Spatial adjacency is recorded enzymatically by copying barcodes of adjacent molecules into a single DNA strand. Because written features exclude DNA, they create gaps in the proximity network. Each reaction product represents one barcode pair, with the reaction’s reach length, the distribution of distances over which pairs can form, determining the spatial resolution of the resulting graph. Reliable reconstruction requires each barcode to connect to several neighbors (> 2, Fig. S1), providing sufficient constraints to infer relative positions. The reaction is independent of the encoded data, and the resulting barcode-pair library constitutes the archival storage medium.

To read the data, the proximity library is sequenced. The collection of sequenced barcode pairs defines a graph: each unique barcode is a node, and each observed pair an edge. Because pairs preferentially connect nearby molecules, the graph contains information about their relative positions. We use the STRND framework (*36*) to reconstruct barcode coordinates, up to rotation, reflection and uniform scale. STRND embeds the graph using random walks, then reduces the high-dimensional embedding to two dimensions through manifold learning. Finally, the reconstructed coordinates are converted into a density image, and a custom image-analysis algorithm identifies barcode-depleted regions (holes) to decode the stored bit pattern.

The feasibility of SPAN storage depends on three physical length scales: the size of the written features (*B*), the mean spacing between DNA molecules (*D*), and proximity interaction range (*L*). Accurate reconstruction requires sufficient graph connectivity (*31*), while resolving written features requires *D* and *L* to be small relative to *B*. We simulated proximity graphs of a 101-bit pattern with 20-µm features and reconstructed them using STRND (Fig. 1B; broader parameter sweep in Fig. S2). At *D*=1µm and *L*=10µm, all 101 bits were recovered without errors. Increasing either parameter introduced high decoding errors. Importantly, these decoding constraints depend on the dimensionless ratios *D*/*B* and *L*/*B*, suggesting that the same principle could extend from the 30-µm features demonstrated below to the 150-nm scale of commercial optical writing, provided molecular spacing and/or interaction range can be reduced accordingly. Writing cost is therefore governed primarily by the patterning technology, while readout cost depends on the number of molecules and proximity pairs required per bit. Efficient storage therefore requires matching the molecular spacing and interaction range to the writing resolution, minimizing the sequencing burden without compromising reconstruction fidelity.

### Demonstration

To experimentally demonstrate SPAN, we encoded a 136-bit (17-byte) pattern representing the MIT logo onto a glass substrate. Using UV photolithography, a 17 × 8 array of 15-nm-thick TiO_2_ features was patterned onto azide-functionalized glass (Fig. 2A). Each feature was 30 µm in diameter, with a pitch of 50 µm, yielding a total patterned area of 830 × 380 µm^2^ (Fig. 2B). The TiO_2_ features define DNA-depleted regions, encoding binary 1, while the surrounding DNA-rich regions encode binary 0.

**Fig. 2.**
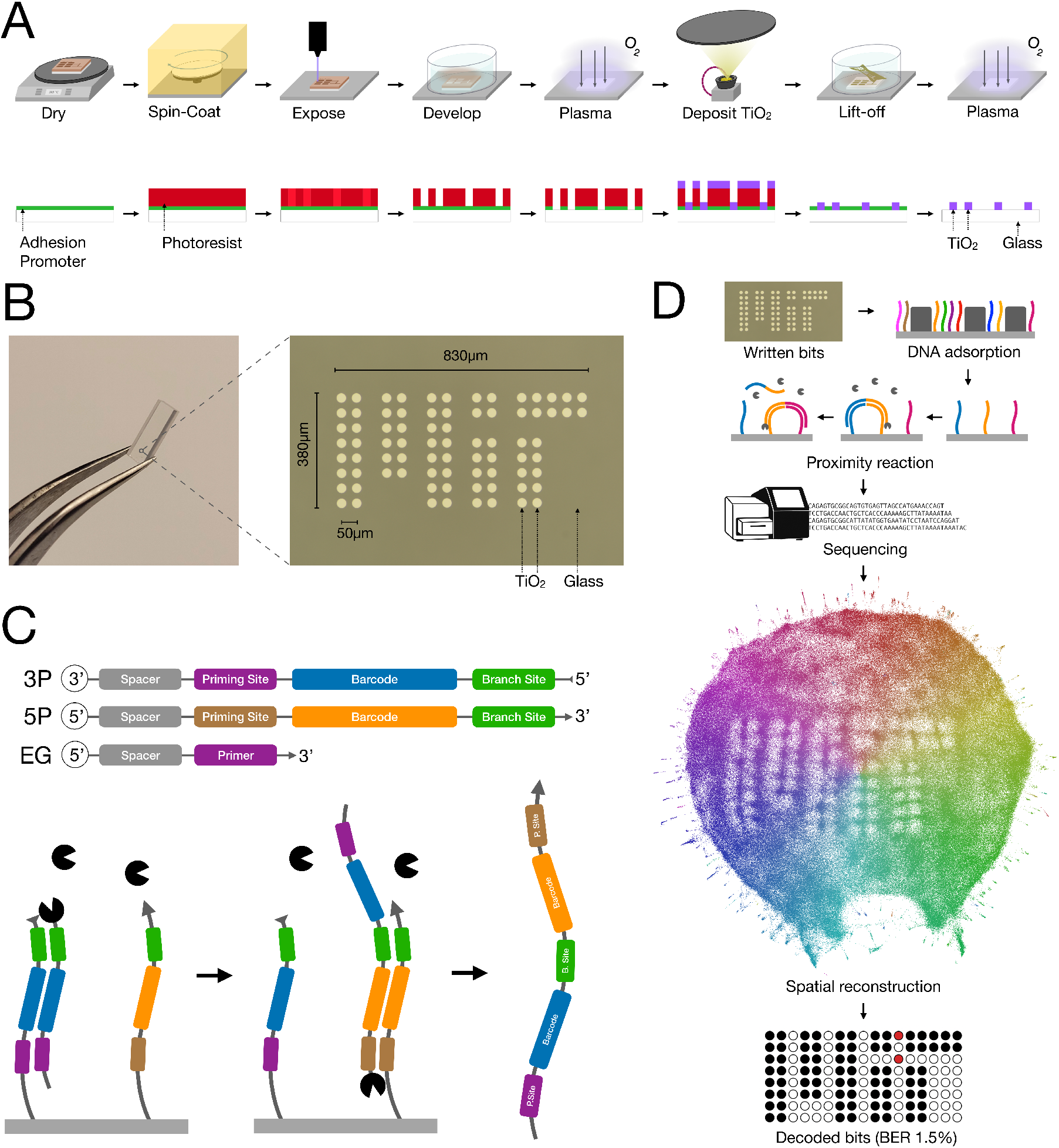
Writing and reading a bit pattern. (**A**) Fabrication of patterned chips. An azide-coated glass slide is coated with photoresist, patterned by UV lithography and developed. Following plasma cleaning, 15nm of TiO_2_ is deposited by electron-beam evaporation. Lift-off leaves TiO_2_ features on glass; a second plasma cleaning removes the remaining azide layer. (**B**) Written bit pattern. Left, a diced 2.5×6 mm chip; right, an optical micrograph showing a 17×8 array encoding the 136-bit (17-byte) MIT logo. Circular features are 30µm in diameter on a 50µm lattice. (**C**) Proximity reaction. The oligo library contains two barcoded oligos (3P and 5P) and an edge generator (EG). Each barcoded oligo contains a unique barcode and a common branch site. EG hybridizes to the priming site of 3P and is extended by DNA polymerase (Pac-Man symbols), copying the first barcode and branch site. Following thermal dissociation, extended EG hybridizes to a nearby 5P oligo via the branch site and is extended to copy the second barcode, generating a double-barcode product. (**D**) Reading the written pattern. After DNA adsorption, enzymatic proximity reaction and sequencing, spatial graph reconstruction reveals DNA-depleted regions as holes in the point cloud. Decoding recovers 134 of 136 bits (bit-error rate, 1.5%). The reconstruction is invariant to rotation and reflection and was manually aligned to the written pattern for display. Microfabrication flow illustrated using Fabublox (*37*).

We next deposited randomly barcoded DNA oligonucleotides (oligos, Fig. 2C) onto the patterned surface. The library comprised two oligo designs, 3P and 5P, each carrying a 25-nt barcode but differing in the position of their priming sites. A third oligo, the edge generator (EG), recorded molecular proximity through a single-pot thermocycled polymerase-extension reaction. EG first hybridizes to the priming site of 3P and is extended to copy its barcode and a shared domain, termed the branch site. Following thermal dissociation, the extended EG hybridizes to the branch site of a nearby 5P, allowing a second extension to copy its barcode and priming site. The resulting products contain two spatially proximal barcodes flanked by priming sites and are amplified by PCR for sequencing.

Next, we sequenced the resulting library using next-generation short-read sequencing (Materials and Methods). Extracted barcode pairs were assembled into a bipartite proximity graph. To reduce noise in the proximity graph, we applied an indirect-path filtering approach adapted from Kloosterman et al. (*38*), retaining edges whose endpoints were also connected by a short alternative path through the graph. The filtered graph contained approximately 613K nodes and 1.1M edges (mean degree 3.7). We then reconstructed the graph in two dimensions using STRND (*36*), assigning spatial coordinates to individual barcodes.

The reconstructed barcode positions form a point cloud in which the written pattern appears as regions of reduced barcode density (Fig. 2D). To decode the pattern, we converted the point cloud into a grayscale density image and fitted a 17×8 lattice using manually selected fiducials. The mean intensity around each lattice site showed a bimodal distribution, allowing threshold-based distinction between DNA-rich (binary 0) and DNA-depleted (binary 1) regions. By selecting 8 fiducial points, we correctly decoded 134 of the 136 written bits, corresponding to a bit-error rate of 1.5%. We note that reconstructions lack an intrinsic physical scale, yet the known dimensions of the written pattern allow us to calibrate node coordinates and edge lengths. For this demonstration, the measured mean molecular spacing was *D* = 1.4 *µ*m, and the characteristic interaction length was *L* = 19 *µ*m, with a feature size of *B* = 30 *µ*m. This proof-of-concept demonstrates the feasibility of recovering lithographically written information from DNA proximity graphs without data-specific DNA synthesis.

### Cost analysis

To assess the economic benefit of SPAN, we compared its writing and readout costs with conventional electronic and DNA storage technologies (Table 1). We consider a scenario in which 1 GB of data is written once and preserved for 100 years, with retrieval performed only when required. For flash storage, we assume a commercial cost of $100 per TB and replacement every ten years, yielding a hardware cost of $1/GB over a century, and negligible readout cost. For DNA storage, we evaluate writing and readout costs separately.

**Table 1.**
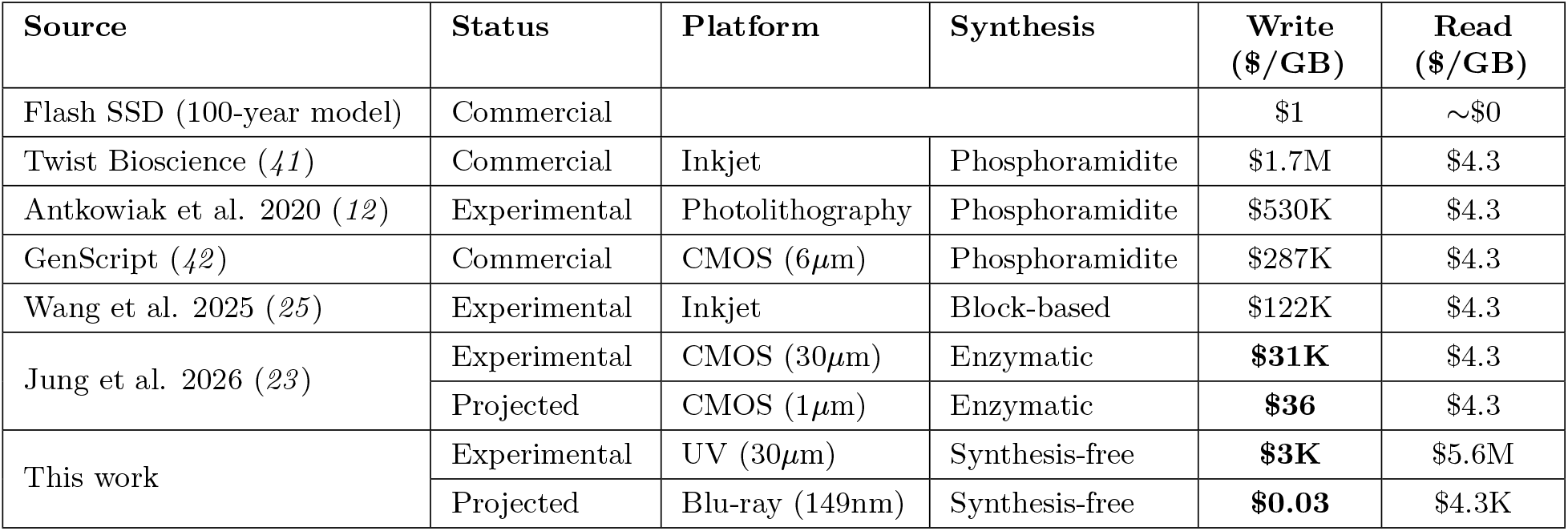
Archival data storage costs over 100 years. Costs are normalized to 1 GB using commercial pricing, published experimental costs, or stated projections. The flash-storage model assumes hardware replacement every ten years and negligible readout costs. Conventional DNA readout costs are estimated using sequencing at $0.8 per gigabase (*39*). Experimental SPAN writing costs are estimated from the exposure times using the Heidelberg MLA-150, amortized over five years, plus reagent costs. Projected SPAN writing costs are estimated using commercially available optical recording technology (Blu-ray). SPAN readout costs are estimated using sequencing at $0.06 per million reads (*40*). Detailed cost calculations and assumptions are provided in Supplementary Text.

| **Source** | **Status** | **Platform** | **Synthesis** | **Write<br>(\$/GB)** | **Read<br>(\$/GB)** |
|---|---|---|---|---|---|
| Flash SSD (100-year model) | Commercial |  |  | \$1 | ∼\$0 |
| Twist Bioscience (*41*) | Commercial | Inkjet | Phosphoramidite | \$1.7M | \$4.3 |
| Antkowiak et al. 2020 (*12*) | Experimental | Photolithography | Phosphoramidite | \$530K | \$4.3 |
| GenScript (*42*) | Commercial | CMOS (6µm) | Phosphoramidite | \$287K | \$4.3 |
| Wang et al. 2025 (*25*) | Experimental | Inkjet | Block-based | \$122K | \$4.3 |
| Jung et al. 2026 (*23*) | Experimental | CMOS (30µm) | Enzymatic | **\$31K** | \$4.3 |
|  | Projected | CMOS (1µm) | Enzymatic | **\$36** | \$4.3 |
| This work | Experimental | UV (30µm) | Synthesis-free | **\$3K** | \$5.6M |
|  | Projected | Blu-ray (149nm) | Synthesis-free | **\$0.03** | \$4.3K |

In conventional DNA storage, writing cost is dominated by data-specific sequence synthesis. Assuming an encoding efficiency of 1.5 bits per base (*10*), commercial phosphoramidite synthesis by Twist Bioscience (inkjet) and GenScript (CMOS) costs approximately $1.7 million and $287,000/GB, respectively. Alternative approaches include assembling prefabricated DNA blocks by enzymatic ligation ($122,000/GB) (*25*). More recently, parallel enzymatic synthesis on CMOS chips has been demonstrated at an estimated $31,000/GB (*23*). Reducing its feature pitch from 30 to 1 µm could lower this to a projected $36/GB, although such miniaturization would require further advances in device fabrication and enzymatic synthesis.

SPAN replaces data-specific DNA synthesis with spatial patterning, decoupling writing costs from base-by-base encoding. Based on the exposure throughput and amortized cost of the Heidelberg MLA-150 used in our demonstration, we estimate a patterning cost of $2,530/GB, plus $440/GB in reagents (Supplementary Text). Using off-the-shelf UV laser writing equipment at the same feature dimensions, the estimated total writing cost for patterning and reagents could be reduced to $570/GB. Significant reductions could be further achieved by using established commercial optical writing technologies. Blu-ray recorders write marks as small as 149 nm (*43*), and Blu-ray-derived optical systems have been demonstrated for direct photolithography (*44, 45*). Assuming comparable throughput at Blu-ray’s information density of 23 Mbit/mm^2^, we project a patterning cost of $0.01/GB. The smaller area per bit also reduces estimated reagent costs to $0.02/GB, yielding a total projected writing cost of $0.03/GB (Supplementary Text).

The low writing cost of SPAN is accompanied by substantially higher readout costs. For conventional sequence-encoded DNA storage, an encoding efficiency of 1.5 bits per base and sequencing at $0.8 per gigabase (*39*) yield an idealized readout cost of $4.3/GB. SPAN instead recovers information from molecular proximity, requiring substantially more bases per bit. Based on the connectivity and filtering requirements of our experimental graph, we estimate that 5 million reads are needed to reconstruct an area capable of encoding approximately 430 bits at the demonstrated resolution (Supplementary Text). At $0.06 per million reads (*40*), this corresponds to an estimated readout cost of $5.6M/GB. Movie S1 illustrates the effect of sequencing depth on graph reconstruction. Readout costs could be reduced by matching molecular spacing (*D*) and interaction range (*L*) to feature size (*B*). In an idealized scaling scenario with 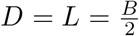, only nine barcode-pair reads per bit are required (Supplementary Text), yielding a projected readout cost of $4,320/GB. Although substantially higher than conventional sequence-based DNA readout costs, these costs could further decline with advances in sequencing technology (*35*).

To assess the economic benefit of SPAN for deep archival storage, we model the expected cost of storing 1 GB of data for 100 years as

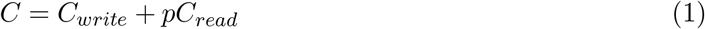

where *C*_*write*_ and *C*_*read*_ are the writing and readout costs (Table 1), and *p* is the probability of retrieval. For projected CMOS-based DNA storage, these costs are $36/GB and $4.3/GB, respectively, compared with $0.03/GB and $4,320/GB for SPAN. As shown in Fig. 3, SPAN becomes less expensive than CMOS-based DNA storage below a read probability of 0.84%. We further compare SPAN with flash memory, modeled at $1/GB over 100 years, including periodic replacement. SPAN’s expected cost falls below flash at read probabilities below 0.02%. These projections suggest a path toward DNA archival storage with lower total expected costs, compared to existing DNA storage systems as well as flash memory, for data that are rarely, if ever, retrieved.

**Fig. 3.**
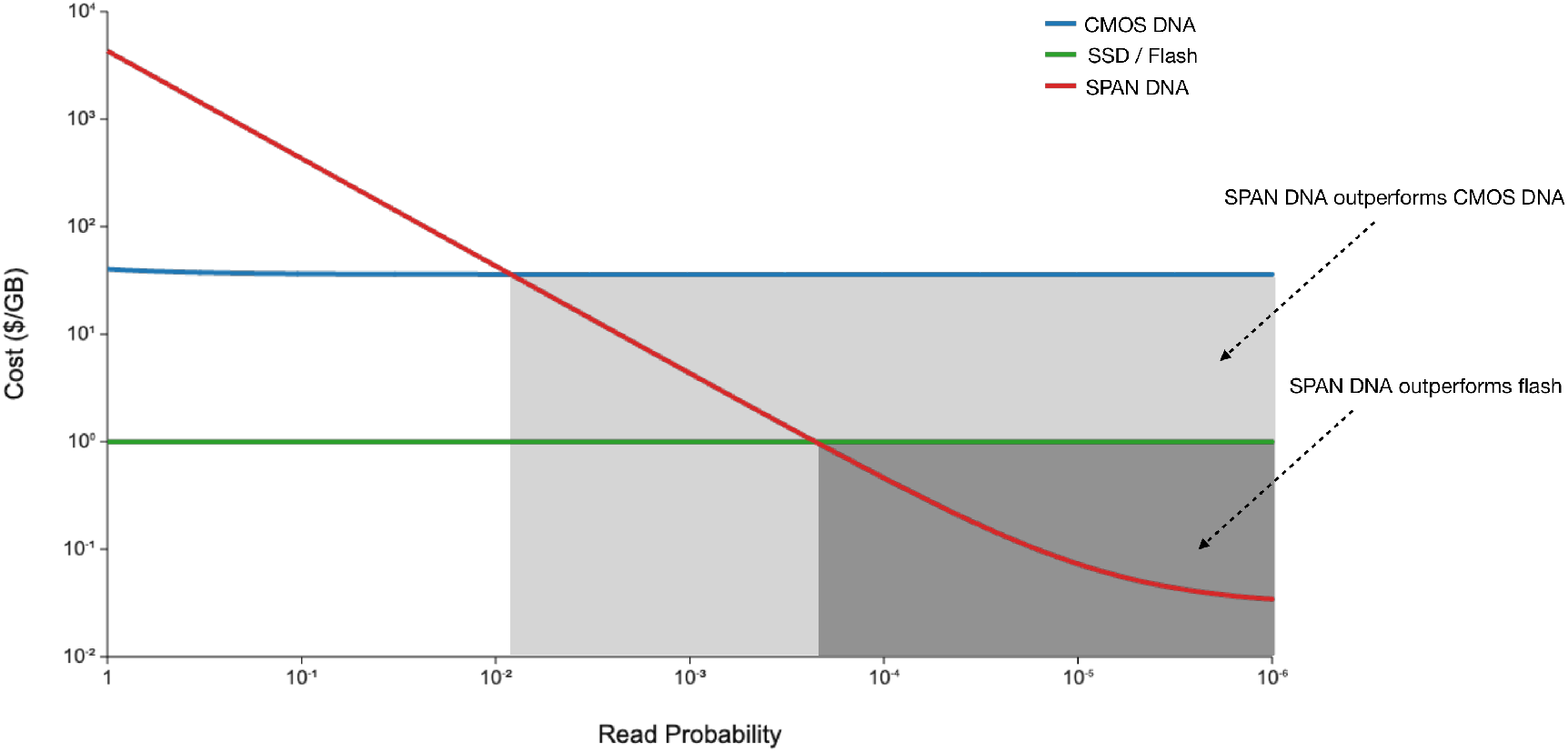
Archival storage cost as a function of read probability. Total expected cost per GB, calculated as *C* = *C*_*write*_ + *pC*_*read*_, for projected CMOS DNA storage and SPAN DNA storage, compared with flash storage modeled at $1/GB over 100 years. SPAN has the lowest expected cost below a read probability of 0.84% compared with CMOS DNA storage, and below 0.02% compared with flash memory.

## Discussion

SPAN introduces a synthesis-independent architecture for molecular data storage, in which information is written as a physical pattern, archived in the proximity relationships of randomly barcoded DNA molecules, and recovered through sequencing and graph reconstruction. We demonstrate this principle by lithographically encoding a 136-bit pattern and recovering 98.5% of the bits. Because neither the barcode sequences nor the enzymatic reactions depend on the information being stored, writing costs are governed primarily by physical patterning and bulk reagents rather than data-specific DNA synthesis. Our cost analysis suggests that this separation could substantially reduce writing costs, at the expense of more demanding readout.

This trade-off is particularly attractive for deep archival storage, where information may be retained for decades or centuries but rarely, if ever, accessed (*2*). Writing and preservation costs may therefore dominate lifetime costs, while retrieval costs are incurred only upon access. Moreover, sequencing costs have declined by several orders of magnitude over the past two decades (*35*), considerably faster than DNA synthesis costs (*6*). If this trend continues, retrieving SPAN archives could become less expensive over time without requiring the data to be rewritten.

Random access (*46*) remains an important challenge for scaling SPAN. Currently, retrieving even a small portion of stored information requires reconstructing the entire proximity graph. One possible solution is spatial indexing: distinct primer sequences (molecular “zip codes”) could be incorporated into the data-independent DNA library and deposited in separate regions using a liquid handler. Upon retrieval, selective PCR amplification would isolate proximity records from the desired region, allowing independent graph reconstruction. Notably, such partitioning would introduce additional dispensing and indexing costs.

Further improvements require tighter control of the three characteristic length scales: written feature size, molecular spacing, and interaction range. Increasing molecular density without reducing feature size increases sequencing demands, while longer interaction ranges improve graph connectivity at the expense of spatial resolution. Efficient storage therefore requires matching molecular spacing and interaction range to the writing resolution. Our thermocycled polymerase-extension reaction provides limited control over interaction distance; alternative proximity-recording approaches (*32, 47*) could enable more localized interactions and reduce sequencing requirements. More sophisticated computational methods for decoding bits from reconstructed graphs could further improve accuracy but remain beyond the scope of this demonstration.

Encoding information in molecular proximity rather than nucleotide sequences may offer further advantages. Because each bit is distributed across multiple molecules, targeted alteration could be more difficult than modifying individual sequences, potentially enabling tamper-evident, write-once archives, although this remains to be explored. Richer per-site alphabets, encoded through feature size or shape, could store multiple bits per site, increasing information density and potentially further reducing writing and reading costs. SPAN could also extend beyond two dimensions: spatially patterned barcode-depleted regions in a three-dimensional medium, such as a hydrogel, might similarly be recovered through proximity-graph reconstruction, potentially increasing volumetric storage density. Recent demonstrations of three-dimensional DNA microscopy (*33*) provide a foundation for such an extension.

SPAN illustrates an alternative approach to molecular information storage in which individual molecules need not be programmed with the information they represent. Instead, information is written as a physical pattern and recovered from the spatial organization of randomly assembled, uniquely identifiable molecules. In this architecture, randomness becomes an asset rather than noise: local molecular interactions reveal the spatial relationships from which the written information can be decoded. This approach suggests that digital information can be stored not only in the sequences of individual molecules, but also in the spatial relationships between them.

## Materials and Methods

### Materials and reagents

Azide-functionalised glass slides were obtained from PolyAn (10400621). Microfabrication used AZ3312 positive photoresist and AZ 300 MIF developer (both MicroChemicals), with isopropanol (IPA) and N-methyl-2-pyrrolidone (NMP) from J.T. Baker. Hydrophobic coating with trichloro-(1H,1H,2H,2H-perfluorooctyl)silane (PFOCTS; Sigma-Aldrich) in BestSolv 7200 (Best Technology).

Molecular biology reagents were Hot Start Taq 2× Master Mix (New England Biolabs, M0496S), 10× PBS (Invitrogen, AM9624), 20× SSC (Thermo Scientific, J60839.K3), 10% SDS (Invitrogen, 15553027), EvaGreen 20X in water (Biotium, 31000-T), and ROX reference dye (Biotium, 29052). All dilutions were prepared in UltraPure DNase/RNase-free distilled water (Invitrogen, 10977023).

Oligonucleotides (oligos) were synthesized by Integrated DNA Technologies at 100–250nmol scale, with machine mixing at degenerate positions. Barcoded oligos were HPLC-purified.

### Bit pattern fabrication

Azide-functionalized glass slides were cleaved into 1×1 inch samples. The azide-functionalized surface, originally designed for DNA immobilization through click chemistry, was used here to promote photoresist adhesion during lithographic patterning. AZ3312 positive photoresist was spin-coated at 3,000 rpm for 45 s and soft-baked at 100 °C for 90 s. Patterns were exposed using a maskless direct-write lithography system (Heidelberg MLA-150), followed by a post-exposure bake at 110 °C for 90 s and development in AZ-300 developer for 90s.

Following development, a 15-nm-thick TiO_2_ layer was deposited by electron-beam evaporation (AJA ATC). Lift-off was performed in an NMP bath at 60 °C with brief sonication, leaving TiO_2_ features at the exposed positions. Samples were subsequently treated in a plasma asher (Thierry Diener) for 5 min at low power to remove the exposed azide coating. Immediately after plasma treatment, samples were immersed in 0.4% (v/v) PFOCTS in BestSolv 7200 for 15 min to produce a hydrophobic surface (*48*). Last, 1×1 inch samples were diced into 2.5×6 mm chips using a dicing saw (Disco DAD-3240), allowing individual chips to be suspended in standard PCR tubes for downstream reactions (a 20uL reaction covers approximately half the surface of a single chip, Fig. S3) .

For the experimental demonstration, a 17×8 binary pattern encoding the MIT logo was fabricated. Individual features were 30µm wide, with a center-to-center pitch of 50 µm, giving a total patterned area of 830×380 µm^2^.

### DNA barcode library

The DNA library comprised three oligonucleotide (oligo) designs: two barcoded oligo (3P and 5P) and an edge-generating oligo (EG). Both barcode-bearing oligo contained a 25-nucleotide variable barcode region, a primer-binding sequence, and a shared hybridization domain. The 3P and 5P designs differed in the relative orientation of their primer-binding sites, positioned toward the 3^′^ and 5^′^ ends, respectively. The EG oligon contained, at its 3^′^ end, a sequence complementary to the primer-binding site of 3P, enabling polymerase extension from a hybridized EG–3P complex.

The available barcode sequence space exceeded 10^13^ distinct sequences, whereas the estimated number of surface-adsorbed barcode molecules was approximately 10^5^-10^6^ per patterned region. The probability of at least one repeated barcode among these molecules was below 5%, as estimated using the birthday-collision approximation

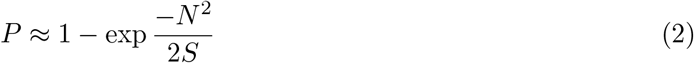

where *N* is the number of molecules and *S* is the barcode sequence space.

For oligo deposition, a solution containing 1µM 3P, 1µM 5P, and 8µM EG in 1X PBS was prepared. A single 2-µL droplet was manually dispensed over the patterned region of each chip. Samples were incubated overnight at room temperature in a closed humidity chamber. Following incubation, chips were washed in 1.5× SSC containing 0.1% SDS for 5 min at 60 °C, followed by 0.5× SSC for 5 min at 60 °C, and rinsed in UltraPure water.

### Proximity reaction

Barcode proximity pairs were generated by a cycled polymerase extension reaction. The EG oligo was designed to hybridize to the primer-binding sequence of a 3P oligo, providing a 3^′^ end for polymerase extension. Extension copies the 3P barcode and the shared hybridization domain, producing an intermediate containing the complement of the first barcode. Following strand dissociation, the extended EG can hybridize through the shared domain to a nearby 5P oligo. A second polymerase extension copies the 5P barcode and its adjacent primer-binding sequence, producing a DNA strand containing both barcode sequences flanked by primer-binding sites.

To generate these products, chips were immersed in 1X Taq Hot Start Master Mix without additional amplification primers. Reactions were thermocycled for 30 cycles, consisting of denaturation at 94 °C for 20s and annealing/extension at 58 °C for 2 min. Repeated hybridization, extension, and dissociation enabled the formation of double-barcode products.

Following the primer-free reaction, a single amplification primer was added to a final concentration of 0.25 µM while the chip remained in the reaction. Additional thermocycling for 20 cycles generated soluble copies of the recorded barcode pairs. The chip was then removed, and the second amplification primer was added to a final concentration of 0.25 µM to enable conventional PCR amplification for 12-18 cycles. Where reaction quantification was required, EvaGreen and ROX reference dye were added during the final amplification step, and product accumulation was monitored by quantitative PCR. The resulting double-stranded products were processed for next-generation sequencing.

### Sequencing

Amplified barcode-pair libraries were sequenced on a Singular G4 platform at the MIT BioMicro Center (Cambridge, MA) using paired-end 50-bp reads, generating up to 100 million reads per sample. A computational pipeline was used to extract the two barcode sequences from each read and identify their corresponding oligo classes (3P or 5P). To correct for sequencing errors, barcodes differing by a Hamming distance of less than 3 were considered equivalent and merged.

### Proximity graph reconstruction

Barcode pairs were assembled into a bipartite graph using the Python package graph-tool (*49*). Each unique barcode was represented as a node, assigned to one of two classes according to its originating oligo design (3P or 5P). An undirected edge was introduced between two nodes when their corresponding barcodes were observed together in a sequencing read, with edge weights representing the number of supporting reads.

To further correct barcode errors, nodes separated by a path of length two on the graph and having a Levenshtein distance of less than 4 were merged. To reduce spurious connections, edges supported by fewer than four reads were discarded, followed by removal of nodes with degree one or degree above 30. We then applied an indirect-path filtering approach adapted from Kloosterman et al. (*38*). For each edge *e* = (*u, v*), we counted the number of alternative paths connecting nodes *u* and *v* without traversing *e*. Because the graph is bipartite, only odd path lengths were considered. We evaluated maximum path lengths of 3, 5, 7, and 9 (Fig. S4). For the MIT pattern, edges were retained only if at least one indirect path of length up to 9 existed between their endpoints.

Spatial coordinates were reconstructed using the STRND framework (*36*). STRND consists of two steps. First, each node is embedded in a high-dimensional space using random walks (*50*), with parameters *d* = 32, *p* = 1, and *q* = 1 (Fig. S5). Second, the high-dimensional embedding is reduced to two dimensions using UMAP (*51*), with 120 nearest neighbors and a minimum distance of 0.5 (Fig. S6). Each barcode is then assigned an (*x, y*) coordinate. The resulting coordinates were exported to CSV as a point cloud for visualization and subsequent bit decoding.

### Point cloud visualization

Reconstructed point clouds were visualized either as red and blue circles according to their oligo class (Figs. S1, S2, S7, and S8; Movie S1), or using a reference-based coloring scheme to compare multiple reconstructions (Fig. 2 and Fig. S4). In the latter approach, multiple reconstructions were generated using different random seeds, and one was selected as a reference. Barcodes were assigned colors from a continuous color wheel according to their positions in the reference reconstruction, and these colors were preserved across all other reconstructions. This approach enabled visual assessment of reconstruction consistency and identification of spatially unstable clusters.

### Decoding bit pattern

To decode individual bits from the reconstructed two-dimensional point cloud, the barcode coordinates were first normalized such that 95% of the points fell within the unit circle. A grayscale density image was then generated by applying a sliding box kernel that counts the number of barcodes within each window. The window size determines the degree of image smoothing; in this study, a window size of 0.5 in normalized units was used (Fig. S7). The resulting density values were mapped to grayscale, with values below the 10th percentile assigned white and values above the 90th percentile assigned black.

To align the written bit pattern with the reconstructed image, 8 fiducial points were manually selected. A regularized quadratic warp (*λ* = 1) was fitted to these fiducials to determine the positions of all 136 lattice sites. For each site, the mean grayscale intensity was calculated within a circle centered on the lattice site. The distribution of mean intensities across the lattice sites was bimodal, allowing the selection of a threshold to distinguish DNA-rich regions (dark, binary 0) from DNA-depleted regions (light, binary 1).

Decoding accuracy was evaluated by comparing the 136 decoded bits with the original written pattern. The bit-error rate (BER) was defined as the fraction of incorrectly decoded bits.

## Acknowledgments

We thank Noah Jakimo for many insightful discussions. This work was carried out in part through the use of MIT.nano’s facilities. We thank the MIT BioMicro Center, and in particular Johan Gibcus, for sequencing services and advice. We acknowledge the MIT Office of Research Computing and Data for providing high-performance computing and data resources that have contributed to the results reported within this paper.

## Author contributions

The research was conceived by J.M.J. and E.P. The microfabrication process was developed by J.O.T. Chip preparation, wet-lab protocol development, data analysis pipeline development, and all experiments and simulations were performed by E.P. The cost analysis was conducted by E.P. and J.M.J. The manuscript was written by E.P. with contributions from all other authors.

## Data, code, and materials availability

Raw sequencing data can be retrieved under 10.6084/m9.figshare.34596324. All code used for data processing, reconstruction, decoding and simulations is available at https://github.com/eyalperry88/SPAN under the MIT license.

## References

1. A. Wright, “Worldwide IDC Global DataSphere Forecast, 2024–2028: AI Everywhere, But Upsurge in Data Will Take Time” (IDC Market Forecast US52076424, International Data Corporation (IDC), 2024).

2. Active Archive Alliance, “Preparing for Tomorrow’s Expanding Storage Challenge with Active Archive” (Annual Report, Active Archive Alliance, 2025; https://activearchive.com/wp-content/uploads/2025/10/AAA-Annual-Report-2025-FINAL.pdf).

3. A. El-Shaikh, B. Seeger, T. Heinis, An Economic Analysis of DNA-based Data Storage Systems. arXiv preprint arXiv:2608.26342 (2026).

4. D. Landsman, “DNA Data Storage Market” in DNA Data Storage: Current Approaches and Emerging Trends, M. K. Gupta, Z. Ping, A. Rasool, B. Cao, P.-Y. Burgi, Eds. (Springer Nature Singapore, Singapore, 2026), pp. 317–332.

5. Intelligence Advanced Research Projects Activity (IARPA), Molecular Information Storage (MIST), https://www.iarpa.gov/research-programs/mist (2019). Office of the Director of National Intelligence.

6. V. Zhirnov, R. M. Zadegan, G. S. Sandhu, G. M. Church, W. L. Hughes, Nucleic acid memory. Nature Materials 15, 366–370 (2016).

7. M. G. Rutten, F. W. Vaandrager, J. A. Elemans, R. J. Nolte, Encoding information into polymers. Nature Reviews Chemistry 2, 365–381 (2018).

8. R. Aksakal, C. Mertens, M. Soete, N. Badi, F. Du Prez, Applications of discrete synthetic macromolecules in life and materials science: Recent and future trends. Advanced science 8, 2004038 (2021).

9. N. Goldman, P. Bertone, S. Chen, C. Dessimoz, E. M. LeProust, B. Sipos, E. Birney, Towards practical, high-capacity, low-maintenance information storage in synthesized DNA. Nature 494, 77–80 (2013).

10. Y. Erlich, D. Zielinski, DNA Fountain enables a robust and efficient storage architecture. Science 355, 950–954 (2017).

11. J. Koch, S. Gantenbein, K. Masania, W. J. Stark, Y. Erlich, R. N. Grass, A DNA-of-things storage architecture to create materials with embedded memory. Nature Biotechnology 38, 39–43 (2020).

12. P. L. Antkowiak, J. Lietard, M. Z. Darestani, M. M. Somoza, W. J. Stark, R. Heckel, R. N. Grass, Low cost DNA data storage using photolithographic synthesis and advanced information reconstruction and error correction. Nature Communications 11, 5345 (2020).

13. D. Bar-Lev, O. Sabary, E. Yaakobi, The zettabyte era is in our DNA. Nature Computational Science 4, 813–817 (2024).

14. G. M. Church, Y. Gao, S. Kosuri, Next-Generation Digital Information Storage in DNA. Science 337, 1628 (2012).

15. L. Ceze, J. Nivala, K. Strauss, Molecular digital data storage using DNA. Nature Reviews Genetics 20, 456–466 (2019).

16. L. Organick, Y.-J. Chen, S. Dumas sAng, R. Lopez, X. Liu, K. Strauss, L. Ceze, Probing the physical limits of reliable DNA data retrieval. Nature Communications 11, 616 (2020).

17. A. L. Gimpel, A. Remschak, W. J. Stark, R. Heckel, R. N. Grass, Comparison of state-of-the-art error-correction coding for sequence-based DNA data storage. Nature Communications 17, 3963 (2026).

18. R. N. Grass, R. Heckel, M. Puddu, D. Paunescu, W. J. Stark, Robust Chemical Preservation of Digital Information on DNA in Silica with Error-Correcting Codes. Angewandte Chemie International Edition 54, 2552–2555 (2015).

19. T. van der Valk, P. Pečnerová, D. Díez-del Molino, A. Bergström, J. Oppenheimer, S. Hartmann, G. Xenikoudakis, J. A. Thomas, M. Dehasque, E. Sağlican, F. R. Fidan, I. Barnes, S. Liu, M. Somel, P. D. Heintzman, P. Nikolskiy, B. Shapiro, P. Skoglund, M. Hofreiter, A. M. Lister, A. Götherström, L. Dalén, Million-year-old DNA sheds light on the genomic history of mammoths. Nature 591, 265–269 (2021).

20. H. H. Lee, R. Kalhor, N. Goela, J. Bolot, G. M. Church, Terminator-free template-independent enzymatic DNA synthesis for digital information storage. Nature Communications 10, 2383 (2019).

21. B. H. Nguyen, C. N. Takahashi, G. Gupta, J. A. Smith, R. Rouse, P. Berndt, S. Yekhanin, D. P. Ward, S. D. Ang, P. Garvan, H.-Y. Parker, R. Carlson, D. Carmean, L. Ceze, K. Strauss, Scaling DNA data storage with nanoscale electrode wells. Science Advances 7, eabi6714 (2021).

22. Z. Weng, J. Li, Y. Wu, X. Xiu, F. Wang, X. Zuo, P. Song, C. Fan, Massively parallel homogeneous amplification of chip-scale DNA for DNA information storage (MPHAC-DIS). Nature Communications 16, 667 (2025).

23. W.-B. Jung, H. S. Jung, J. Wang, H. Hinton, S. J. Kim, Y. Zhang, S. Chen, Y.-H. Hwang, M. Fournier, M. Boul, K. Grosselin, A. Horgan, X. Godron, R. Nicol, D. Ham, Parallel enzymatic DNA synthesis using a semiconductor chip. Nature Electronics 9, 932–940 (2026).

24. S. Jo, H. Shin, S.-y. Joe, D. Baek, C. Park, H. Chun, Recent progress in DNA data storage based on high-throughput DNA synthesis. Biomedical Engineering Letters 14, 993–1009 (2024).

25. C. Wang, D. Wei, Z. Wei, D. Yang, J. Xing, Y. Wang, X. Wang, P. Wang, G. Ma, X. Zhang, H. Li, C. Tang, P. Hou, J. Wang, R. Gao, G. Xie, C. Li, Y. Ju, P. Wang, L. Yue, Y. Zhao, Y. Sheng, J. Xiao, H. Niu, S. Xu, H. Yang, D. Liu, B. Duan, D. Bu, G. Tan, F. Chen, Cost-Effective DNA Storage System with DNA Movable Type. Advanced Science 12, 2411354 (2025).

26. C. Zhang, R. Wu, F. Sun, Y. Lin, Y. Liang, J. Teng, N. Liu, Q. Ouyang, L. Qian, H. Yan, Parallel molecular data storage by printing epigenetic bits on DNA. Nature 634, 824–832 (2024).

27. E. N. Glezer, M. Milosavljevic, L. Huang, R. J. Finlay, T.-H. Her, J. P. Callan, E. Mazur, Three-dimensional optical storage inside transparent materials. Optics Letters 21, 2023–2025 (1996).

28. Microsoft Research Project Silica Team, Laser writing in glass for dense, fast and efficient archival data storage. Nature 650, 606–612 (2026).

29. E. Lieberman-Aiden, N. L. Van Berkum, L. Williams, M. Imakaev, T. Ragoczy, A. Telling, Amit, B. R. Lajoie, P. J. Sabo, M. O. Dorschner, et al., Comprehensive mapping of long-range interactions reveals folding principles of the human genome. science 326, 289–293 (2009).

30. I. T. Hoffecker, Y. Yang, G. Bernardinelli, P. Orponen, B. Högberg, A computational framework for DNA sequencing microscopy. Proceedings of the National Academy of Sciences 116, 19282–19287 (2019).

31. J. A. Weinstein, A. Regev, F. Zhang, DNA Microscopy: Optics-free Spatio-genetic Imaging by a Stand-Alone Chemical Reaction. Cell 178, 229-241.e16 (2019).

32. F. Karlsson, M. Simonetti, C. Galonska, M. Karlsson, H. van Ooijen, D. Thiagarajan, T. Kallas, M. Schweitzer, L. Larsson, V. van Hoef, P. Tajvar, J. Dahlberg, F. De Temmerman, L. Leijo-nancker, V. Trombin, S. Geny, R. Forlin, E. Negrini, B. Wu, L. Xi, S. Petkov, L. Franzén, J. Bunz, C. Moge, H. Everberg, P. Brodin, A. Martinez Barrio, S. Fredriksson, Cells as Single-Molecule Protein Proximity Networks. bioRxiv, doi: 10.1101/2025.06.19.660329 (2025). Preprint.

33. N. Qian, J. A. Weinstein, Spatial transcriptomic imaging of an intact organism using volumetric DNA microscopy. Nature Biotechnology 44, 222–232 (2026).

34. H. Liao, S. Kottapalli, Y. Huang, M. Chaw, J. Gehring, O. Waltner, M. Phung-Rojas, R. M. Daza, F. A. Matsen, C. Trapnell, J. Shendure, S. Srivatsan, Optics-free reconstruction of shapes, images and volumes with DNA barcode proximity graphs. eLife 15, RP110436 (2026). eviewed Preprint (version 2).

35. K. A. Wetterstrand, DNA Sequencing Costs: Data from the NHGRI Genome Sequencing Program (GSP), National Human Genome Research Institute (2023); https://www.genome.gov/about-genomics/fact-sheets/DNA-Sequencing-Costs-Data. Last updated May 16, 2023.

36. D. Fernandez Bonet, I. T. Hoffecker, Image recovery from unknown network mechanisms for DNA sequencing-based microscopy. Nanoscale 15, 8153–8157 (2023).

37. Fabublox,; https://fabublox.com/. SPAN Process flow, to be published.

38. A. Kloosterman, I. Baars, B. Högberg, An error correction strategy for image reconstruction by DNA sequencing microscopy. Nature Computational Science 4, 119–127 (2024).

39. Ultima Genomics, Whole genome sequencing cost, Web page; https://www.ultimagenomics.com/focus-areas/whole-genome-sequencing/. Accessed October 11, 2026.

40. Roche, Roche Diagnostics Day 2026, Investor presentation, slide 98 (2026); https://assets.roche.com/f/176343/x/d7d2be3148/diagnositcs-day-2026.pdf. Presented May 12, 2026.

41. Twist Bioscience, Twist oligo pools, Product sheet; https://www.twistbioscience.com/resources/product-sheet/twist-oligo-pools. Accessed October 11, 2026.

42. GenScript, Oligo pools, Product specifications; https://www.genscript.com/oligo-pools.html. Accessed October 11, 2026.

43. ISO/IEC, “30190:2021” (ISO/IEC, 2021; https://www.iso.org/standard/80770.html). 3rd edition. Single-layer (25 GB) disk specifications.

44. C. A. Rothenbach, M. C. Gupta, High resolution, low cost laser lithography using a Blu-ray optical head assembly. Optics and Lasers in Engineering 50, 900–904 (2012).

45. P.-Y. Chen, W.-Y. Jywe, M.-S. Wang, C.-H. Wu, Application of blue laser direct-writing equipment for manufacturing of periodic and aperiodic nanostructure patterns. Precision Engineering 46, 263–269 (2016).

46. L. Organick, S. D. Ang, Y.-J. Chen, R. Lopez, S. Yekhanin, K. Makarychev, M. Z. Racz, G. Kamath, P. Gopalan, B. Nguyen, C. N. Takahashi, S. Newman, H.-Y. Parker, C. Rashtchian, K. Stewart, G. Gupta, R. Carlson, J. Mulligan, D. Carmean, G. Seelig, L. Ceze, K. Strauss, Random access in large-scale DNA data storage. Nature Biotechnology 36, 242–248 (2018).

47. T. E. Schaus, S. Woo, F. Xuan, X. Chen, P. Yin, A DNA nanoscope via auto-cycling proximity recording. Nature Communications 8, 696 (2017).

48. A. Toppi, M. Dufva, Accessible, fast and easy fabrication of hydrophilic-in-hydrophobic micro-droplet arrays. PLOS ONE 17, e0263282 (2022).

49. T. P. Peixoto, The graph-tool python library. figshare, doi: 10.6084/m9.figshare.1164194 (2014).

50. A. Grover, J. Leskovec, “node2vec: Scalable feature learning for networks” in Proceedings of the 22nd ACM SIGKDD international conference on Knowledge discovery and data mining (2016), pp. 855–864.

51. L. McInnes, J. Healy, J. Melville, Umap: Uniform manifold approximation and projection for dimension reduction. arXiv preprint arXiv:1802.03426 (2018).

52. Heidelberg Instruments, MLA 150 maskless aligner, Product specifications; https://heidelberg-instruments.com/product/mla150/. Accessed October 11, 2026.

53. Pohang University of Science and Technology (POSTECH), Maskless Laser Lithography (MLA150), Research Equipment Support Portal (RETINA), equipment record no. 10152128 (n.d.); https://rtsc.postech.ac.kr/center/equipment/view.jsp?eqid=10152128. Acquired April 3, 2020, for KRW 569,860,860; accessed October 31, 2026.

54. xTool, xTool F2 Ultra UV User Guide (2026); https://support.xtool.com/article/2359. Updated October 26, 2026.

55. Kioxia Corporation, Sandisk Corporation, New 3D Flash Memory Technology from Kioxia and Sandisk Achieves Industry’s Highest Bit Density for QLC NAND (2026); https://apac.kioxia.com/en-apac/about/news/2026/20260804-2.html. Published August 4, 2026.

56. Fujifilm North America Corporation, LTO Ultrium 10 (40 TB) Data Cartridge (2026); https://asset.fujifilm.com/www/us/files/2026-04/f504b921d82ca915758da2ea1b660c81/FNAC-DSS_LTO-10_40_TB_Tape_Spec_Sheet-02.2026.pdf. Product specification sheet.

57. Sony Corporation, Panasonic Corporation, “White Paper: Archival Disc Technology” (Sony Corporation and Panasonic Corporation, 2020; https://panasonic.cn/wp-content/uploads/2020/05/Archival-Disc-Technology-%EF%BC%9A2nd-Edition.pdf).

