## Supplementary Information for "Archival data storage using DNA proximity graphs"

**This file includes:**

Supplementary Text
Figs. S1 to S8
Tables S1 to S2

**Other Supplementary Materials for this manuscript include the following:**

Movie S1

**Supplementary Text**

Experimental writing cost estimate

We estimated the cost of writing SPAN at the geometry of the experimental 136-bit (17-byte) demonstration. The values below are extrapolated to 1 GB ($10^9$ bytes) using the demonstrated pattern dimensions and the direct-write photolithography throughput.

*Patterning cost*: The 17-byte pattern occupied 830×380 µm², or 0.3154mm², and was fabricated with a Heidelberg MLA-150 maskless lithography system (*52*). Using a maximum exposure throughput of 1,400 mm²/min gives an exposure time of 13.5ms per pattern. We assume a capital cost of $500K (*53*), amortized over five years of continuous operation (157 million seconds), corresponding to an equipment cost of approximately $3.18 per millisecond. The resulting equipment cost is approximately $0.000043 per 17-byte pattern, equivalent to $2,530/GB.

*Enzymatic reagent cost:* For the experimental patterned area, while we used a 20uL reaction in a PCR tube, we estimate that the reagent volume consumed was localized to not more than the 10µm above the chip surface, comprising of a reaction volume of 3.2nL. At the same information density, writing 1GB requires approximately 185mL of reaction mixture. We use the quoted price of Hot Start Taq 2× Master Mix (New England Biolabs), $274 for 500 reactions of 50µL each, equivalent to $10.96/mL. This gives approximately $2,000/GB at the quoted price. Assuming a tenfold reduction in reagent price at industrial scale, the estimated enzyme cost is approximately $200/GB.

*Oligonucleotide (oligo) cost:* The data-independent library contains three oligo designs (3P, 5P, and EG). Using a length of 60nt per design and a 10µmol-scale synthesis price of $21.50 per nucleotide from Integrated DNA Technologies, each design costs $1,290, or $3,870 for all three. At the experimental concentrations of 1µM each for 3P and 5P and 8µM for EG, 185mL of reaction volume consumes approximately 185nmol of each barcode oligo and 1.4µmol of EG. Allocating the bulk synthesis prices proportionally to consumption gives an oligo cost of approximately $240/GB.

*Combined cost:* Combining $2,530/GB for patterning with $200/GB for enzymes and $240/GB for oligos gives approximately **$3,000/GB**

Off-the-shelf writing cost estimate

Next, we estimate the writing cost of SPAN using commercially available UV laser equipment while retaining the same geometry and information density as the experimental demonstration (50µm pitch, 30µm features). As a commercial reference, the xTool F2 Ultra UV laser engraver (*54*) uses a 355nm laser, supports scanning speeds up to 15,000 mm/s and pulse frequencies up to 150kHz, and costs approximately $4,000.

*Patterning cost:* We assume a capital cost of $4,000, amortized over five years of continuous operation (157 million seconds), corresponding to $2.5/µs. At a pulse frequency of 150 kHz, we assume a scanning speed of 750 mm/s, giving 5µm spacing between pulses. Raster exposure of the entire 830×380 $\mu m^2$ pattern with 5μm line spacing requires 76 passes, totaling 63.1mm of scanning distance. The resulting exposure time is 84ms per 17-byte pattern, corresponding to an equipment cost of approximately $130/GB.

*Reagent and oligo costs:* Since the geometry and information density remain unchanged, we retain the estimates above of $200/GB for enzymes and $240/GB for oligos.

*Combined cost:* Combining $120/GB for patterning with $200/GB for enzymes and $240/GB for oligos gives approximately **$570/GB**.

Projected writing cost estimate

To estimate the potential writing cost of SPAN at scale, we use Blu-ray optical recording as a benchmark for mature, inexpensive, high-density patterning. Blu-ray systems use a 405nm laser and a high-numerical-aperture objective (NA=0.85), enabling recorded marks as short as 149nm. Consumer Blu-ray recorders operating at 8× speed can write 25GB in approximately 12 minutes. Photolithography using optical equipment derived from Blu-ray recorders has also been demonstrated (*44, 45*). We therefore consider a custom Blu-ray-derived optical system adapted to pattern DNA-compatible substrates.

*Patterning cost:* We assume a custom optical writing system costing $50,000, amortized over five years of continuous operation. At 12min per 25-GB disc, this corresponds to approximately 219,000 discs over the equipment lifetime. The resulting estimated patterning cost is $0.0091/GB, rounded to $0.01/GB. For subsequent estimates we use an effective areal bit density of 22.8 $Mbit/mm^2$, equivalent to approximately 209nm spacing on a square lattice.

*Enzymatic reagent cost:* Reagent consumption is estimated as above. Reducing the bit pitch from 50μm to approximately 209nm raises areal density by a factor of X57,000, lowering the estimated reaction volume from about 185mL/GB to about 3.2μL/GB. At the assumed industrial enzyme price, this corresponds to approximately $0.0035/GB. We round this component upward to $0.01/GB.

*Oligo cost:* SPAN uses the same data-independent oligo library irrespective of the written dataset. We retain the commercial synthesis price and oligo lengths assumed above. Conservatively assuming a 5μL volume for the projected 1GB pattern, at concentrations of 1μM each for 3P and 5P and 8μM for EG, the reaction consumes 5pmol each of 3P and 5P and 40 pmol of EG. Allocating the synthesis cost to these quantities gives approximately $0.0065/GB, rounded upward to $0.01/GB.

*Combined cost:* Combining $0.01/GB for patterning with $0.01/GB for enzymes and $0.01/GB for oligos gives approximately **$0.03/GB**.

Experimental reading cost estimate

To estimate the readout cost of our experimental demonstration, we consider the graph obtained after filtering noisy nodes and edges, as described in Materials and Methods. This graph contains 612,719 nodes. Based on our reconstruction results, we assume that a mean node degree of three is required for a sufficiently connected graph to achieve stable reconstruction (Fig. S1). This corresponds to approximately 919,000 unique edges. As our filtering procedure requires each retained edge to be supported by at least four sequencing reads, a minimum of 3.7 million reads is required. Allowing for 25% of reads to be uninformative, for example due to leaf nodes or discarded edges, we estimate that approximately 5M reads are needed to reconstruct the graph.

Although the reconstructed coordinates lack an intrinsic physical scale, the known dimensions of the MIT pattern allow us to estimate the area covered by the reconstructed graph. The reconstructed region spans a circle approximately 1.2 mm in diameter, which could accommodate ~430 bits using the same feature diameter (30 µm) and pitch (50 µm) as our experimental demonstration, with all features contained within the circle. Thus, we estimate that 5M sequencing reads are required to decode 430 bits. At the reported sequencing cost of $0.06 per million reads on Roche's AXELIOS 1 platform (*40*), this corresponds to $0.3 for 430 bits. Scaling to 1GB, the estimated readout cost is approximately **$5.6M/GB**.

Minimal number of reads per bit

To estimate the potential reading cost of SPAN at scale, we are interested in the question, what is the minimum number of barcode pairs required to decode a single bit. We consider a honeycomb lattice as an ideal 2D molecular proximity graph. In this arrangement, the molecular spacing $D$ and interaction reach $L$ are both equal to some distance $d$, such that every barcode has exactly three neighbors.

We first consider the minimum diameter $B$ of a circular written feature that guarantees intersection with at least one barcode, regardless of its position. If a written feature is smaller than the spacing between the barcodes, its existence or lack of cannot be decoded in any graph reconstruction. In a honeycomb lattice, the largest distance between any point and its nearest node is $d$, occurring at the center of a hexagonal cell. Therefore, the minimum diameter required to guarantee intersection with at least one node is $B=2d$.

Thus, we set the written feature length of $B=2d$. These written features can be positioned to overlap exactly two adjacent barcodes (the distance between any two adjacent nodes on our graph is exactly $d$). Having a written feature on the lattice effectively removes the pair of nodes and their associated edges from the graph. We therefore ask how many such pairs, $P$, representing $P$ encoded bits, can be removed while preserving a single connected graph spanning the original lattice. The pairs have to be isolated. No pair can share a node or a neighbor, for it to be decoded as a stable hole in the reconstructed graph.

For a honeycomb lattice containing $N$ nodes, each of degree 3, the total number of edges is

$$E = \frac{3N}{2}$$

Removing $P$ isolated pairs of adjacent nodes reduces the number of nodes to

$$N' = N - 2P$$

Each removed pair eliminates five edges: one connecting the two nodes and four connecting them to the surrounding lattice. Following our assumption that pairs are well-separated, no edges are shared. Thus, the number of edges in the reduced graph is

$$E' = E - 5P$$

For the reduced graph to be connected, it must satisfy:

$$E' \geq N' - 1$$

We plug the previous equations in:

$$E - 5P \geq N - 2P - 1$$

Plugging in $E = \frac{3N}{2} \rightarrow N = \frac{2E}{3}$, and dropping the constant term, we obtain

$$E \geq 9P$$

Thus, in the ideal honeycomb lattice, at least nine edges are required per bit encoded as a stable hole. A stable hole is defined as a removed pair that does not disrupt graph connectivity. This represents an optimistic yet feasible estimate that we use to calculate the projected number of sequencing reads needed per bit.

Projected reading cost estimate

To estimate the projected readout cost, we use the theoretical minimum of nine barcode-pair reads per bit derived above. Using the reported sequencing cost of \$0.06 per million reads on Roche's AXELIOS 1 platform (*40*), this corresponds to \$0.54 per million bits (Mbit). Scaling to 1GB yields a projected readout cost of approximately \$4,320/GB.

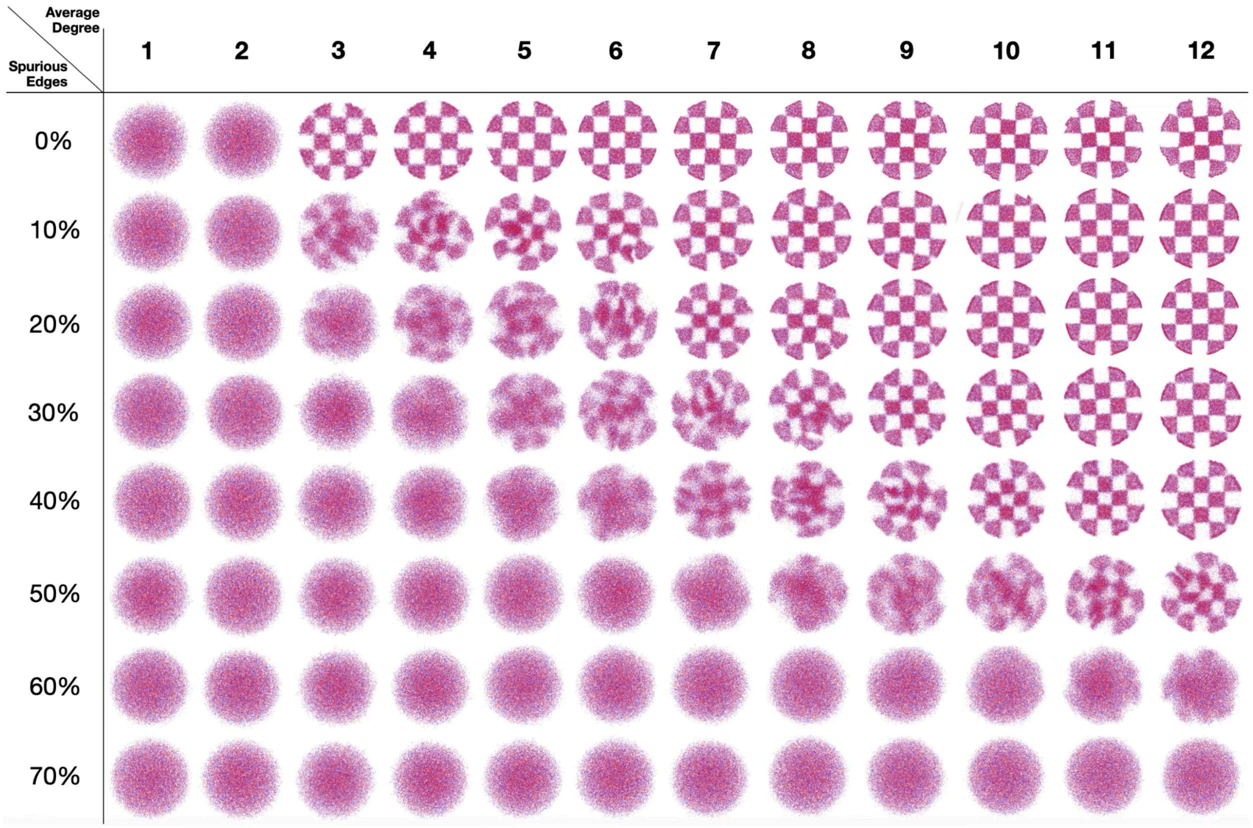


**Fig. S1: Effect of graph connectivity and spurious edges on spatial reconstruction.** Simulated proximity graphs were generated from nodes with known spatial coordinates. True edges connected nodes within a fixed distance, while spurious edges connected randomly selected nodes regardless of distance. Edges were sampled until the specified average degree (columns) and fraction of spurious edges (rows) were reached. The original coordinates were then hidden, and each graph was spatially reconstructed using only its connectivity. Each panel shows the resulting node positions, with the checkerboard pattern indicating reconstruction resolution. Without spurious edges, an average degree of three is sufficient for accurate reconstruction. As the fraction of spurious edges increases, higher connectivity is required, until reconstruction fails at high noise levels.

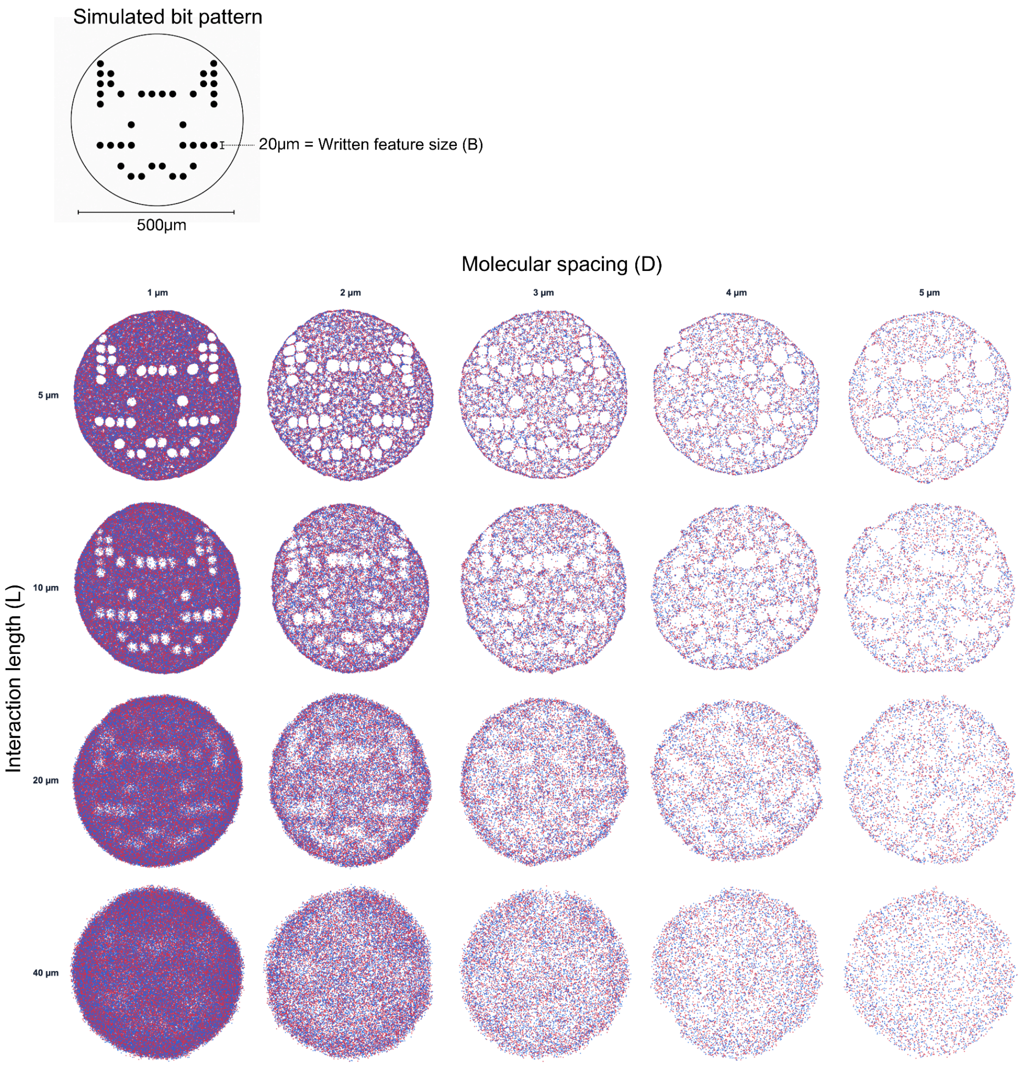


**Fig. S2: Effects of molecular spacing and interaction length on the reconstruction of a bit pattern.** A 12×12 (144-bit) pattern depicting a cat, adapted from the original emoji set designed by Shigetaka Kurita in 1999, is used as ground truth. Each bit is represented by a circular feature of diameter B=20µm, fixed throughout the simulations, arranged on a 30µm center-to-center lattice within a 500µm canvas. Nodes are randomly distributed according to a spatial Poisson process, with density determined by the mean molecular spacing (D), excluding regions corresponding to 1s on the bit pattern. Edges are then sampled with a characteristic interaction length (L) until an average node degree of four is reached, yielding a giant connected component. The resulting graph is reconstructed and visualized as a point cloud, with red and blue dots representing the two node types. When both D and L are sufficiently small relative to B, the original bit pattern is clearly resolved. At short interaction lengths, patterns can also emerge from sparsely sampled graphs, although neighboring features may merge. We note that reconstruction is governed by the dimensionless ratios D/B and L/B, rather than by the absolute physical dimensions of the system.

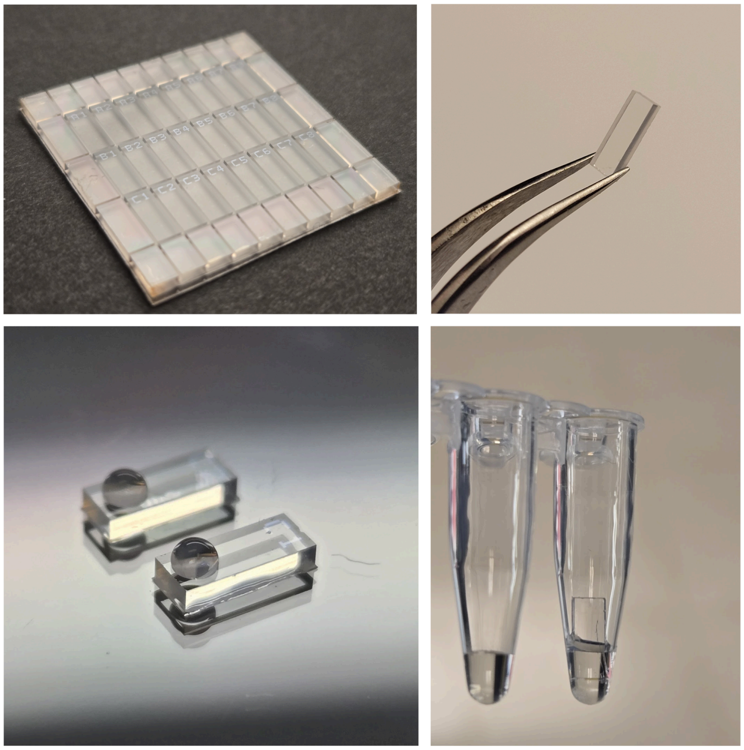

**Fig. S3: Photographs of fabricated chips and experimental setup.** *Top left:* A batch of chips following the complete microfabrication process. A 1×1 inch glass slide is diced into 2.5×6mm chips. Each chip contains a label patterned by deposition of 15nm $TiO_2$, visible to the naked eye. *Top right:* A single 2.5×6 mm chip. *Bottom left:* DNA immobilization by droplet deposition. Two 2µL droplets are deposited onto separate chips. The hydrophobic surface produces high contact angles and prevents droplet spreading. When placed in a humidity chamber, the droplets remain stable overnight. *Bottom right:* A chip immersed in a PCR tube containing 20µL of reaction solution. The solution covers approximately half of the chip, which can be retrieved using tweezers.

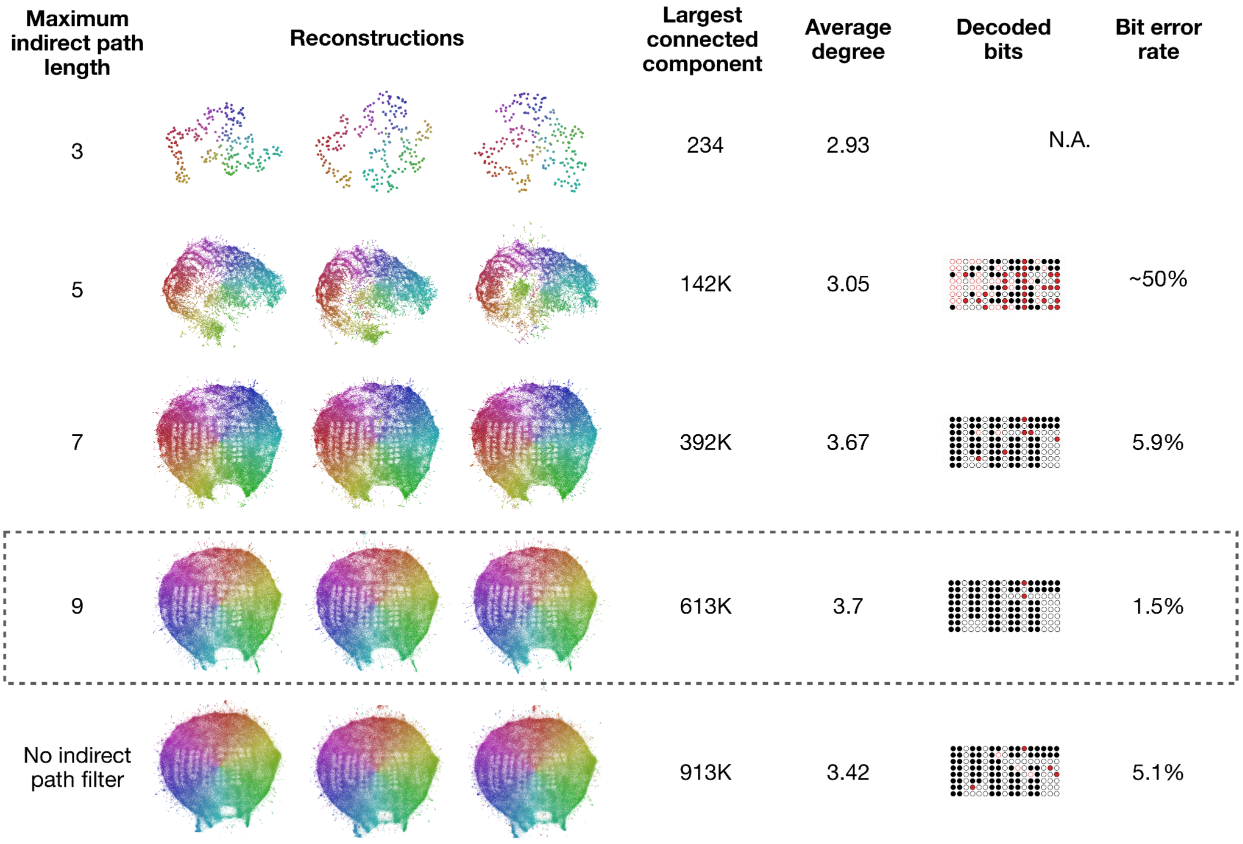


**Fig. S4: Effect of indirect-path filtering on graph reconstruction and decoding accuracy.** Comparison of proximity-graph reconstructions using an indirect-path filtering approach adapted from Kloosterman et al. (*38*), retaining only edges supported by at least one indirect path with a maximum length of 3, 5, 7, or 9, alongside an unfiltered control (bottom row). Each row shows three independent reconstructions using different random seeds, with barcodes colored to visualize reconstruction stability. The number of nodes in the largest connected component, average node degree, decoded bit maps, and bit-error rates are shown for each condition. Using a maximum path length of 3, as originally demonstrated by Kloosterman et al., results in a highly fragmented graph with only 234 nodes in its largest connected component, preventing decoding. Increasing the maximum path length preserves more connected nodes and improves reconstruction quality, with a minimum bit-error rate of 1.5% at a maximum path length of 9, compared with 5.1% without indirect-path filtering. The selected filtering condition used in the main text is highlighted by the dashed outline.

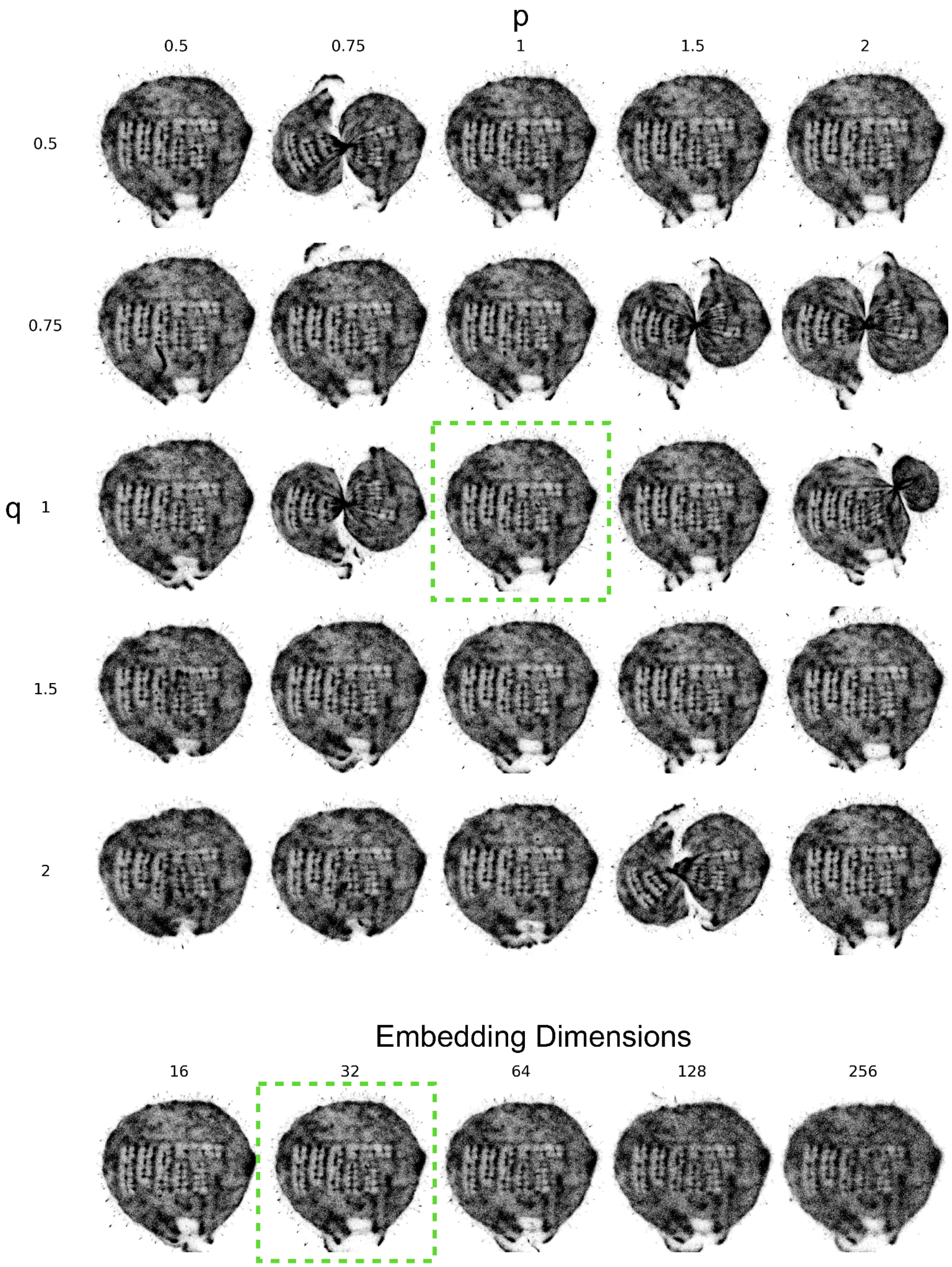


**Fig. S5: Reconstruction Parameter Sweep (Node2Vec).** The matrix shows reconstructions using different parameters for the first stage of STRND (*36*), in which nodes are embedded in a high-dimensional space (Node2Vec (*50*)) using random walks. The parameters p ("return") and q ("in-out") control the random walks, biasing them toward breadth-first search (BFS) or depth-first search (DFS). The embedding dimension (d) determines the number of dimensions used in the resulting embeddings. The parameters used in this study were q=1, p=1, and d=32, similar to those used in the original STRND paper.

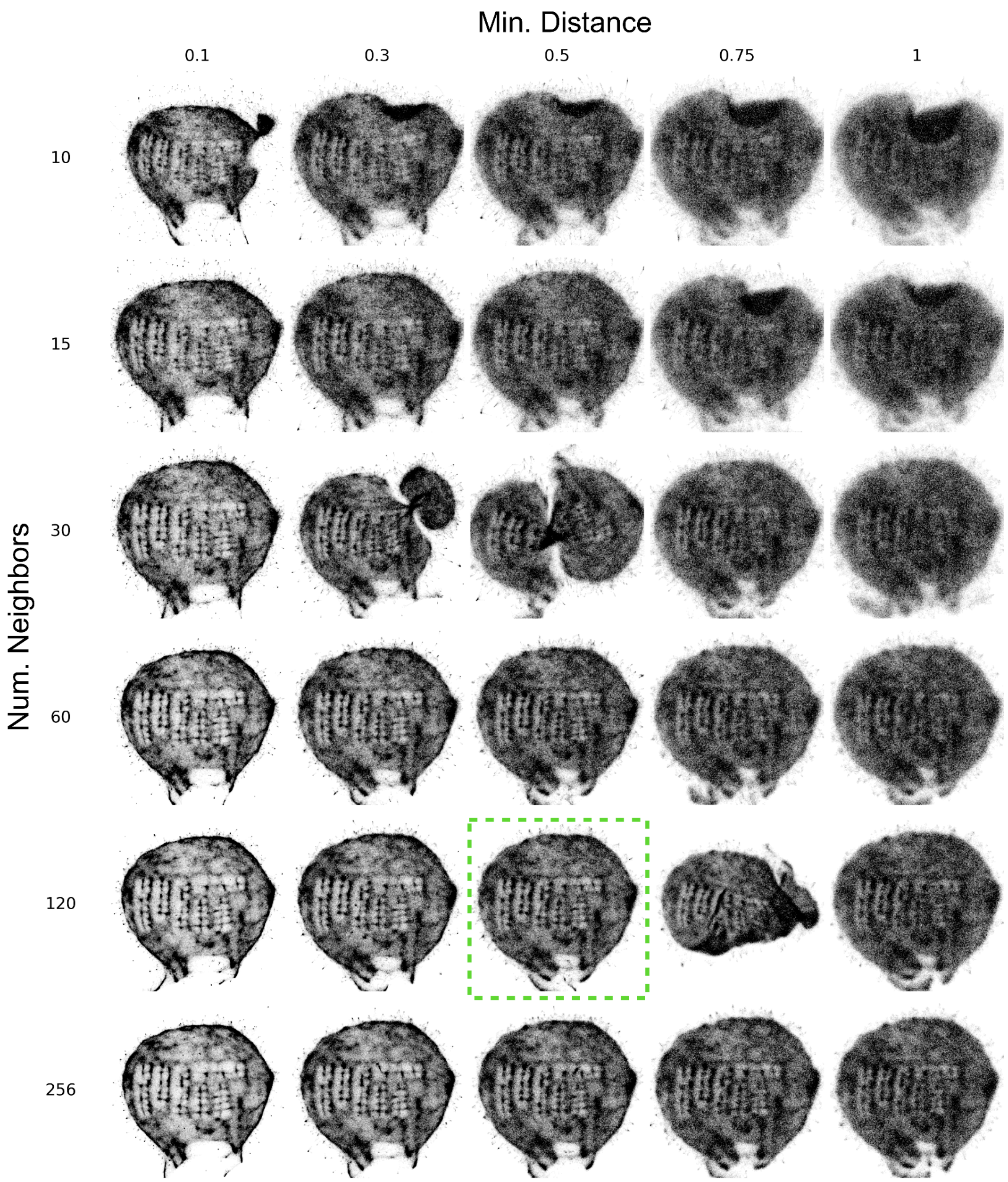


**Fig. S6: Reconstruction Parameter Sweep (UMAP).** The matrix shows reconstructions using different parameters for the second stage of STRND (*36*), in which the high-dimensional embeddings of nodes are reduced to a two-dimensional representation using UMAP (*51*). UMAP uses two main parameters: the number of neighbors (n_neighbors), which controls the balance between preserving local and global structure, with larger values favoring global structure; and the minimum distance (min_dist), which controls how tightly points are packed in the resulting embedding, with smaller values allowing denser clustering. The parameters used in this study were n_neighbors=120 and min_dist=0.5.

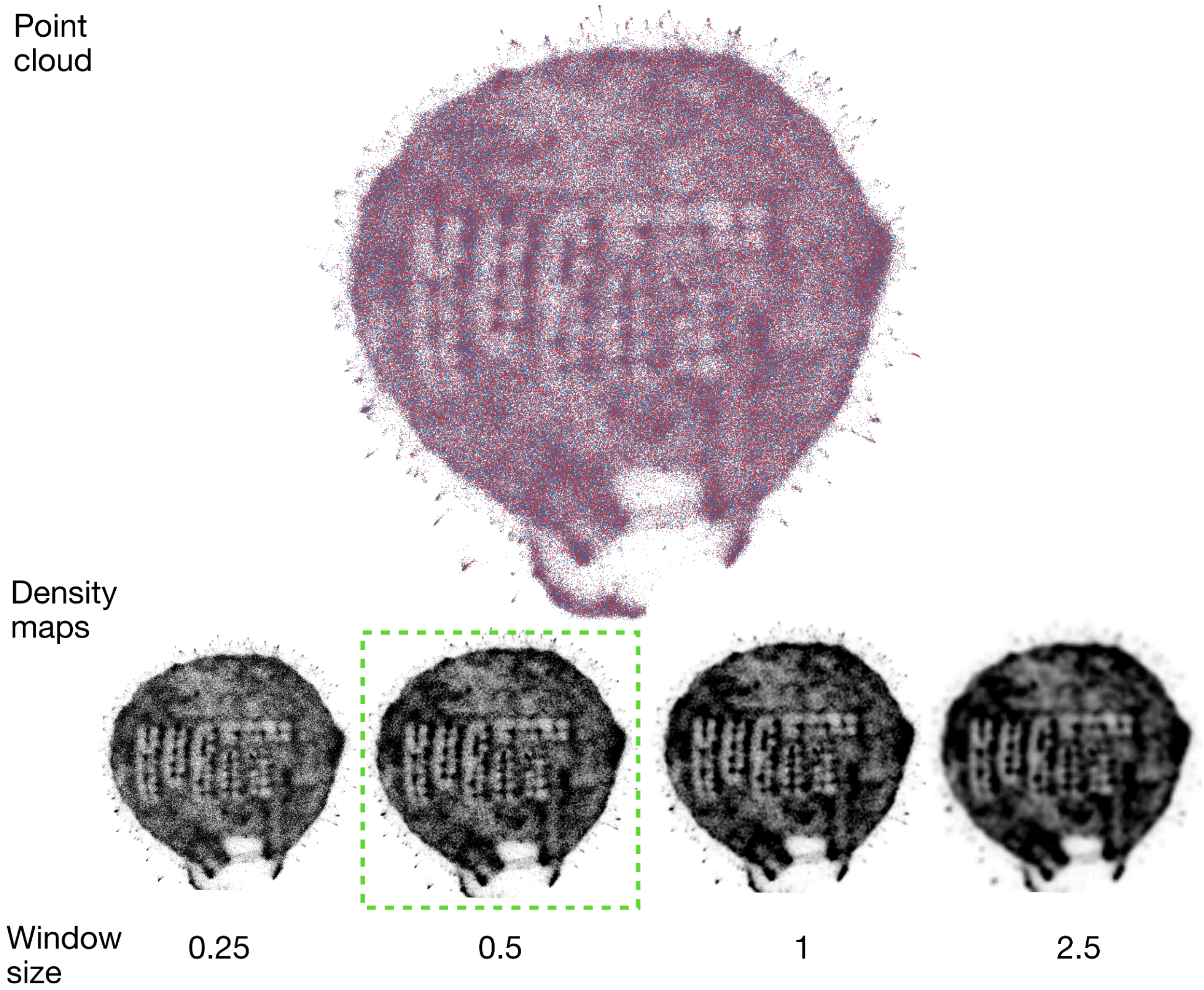


**Fig. S7: From point cloud to density maps.** Reconstruction assigns a 3D coordinate to each barcode, shown as a point cloud at the top of the figure, where red and blue points correspond to 3P and 5P oligos, respectively. For bit decoding, we disregard the distinction between the two classes of oligos and apply a sliding square box kernel to the reconstructed points to generate a density map. First, the coordinates are normalized such that 95% of the points lie within the unit circle. We then define a window size and count the number of barcodes within each window. The resulting density counts are normalized such that values at or below the 10th percentile appear white, and values at or above the 90th percentile appear black, with linear scaling in between. The bottom of the figure shows four density maps generated using different window sizes. Larger windows produce smoother images but can obscure fine features. In this study, we used a window size of 0.5% of the unit-circle diameter.

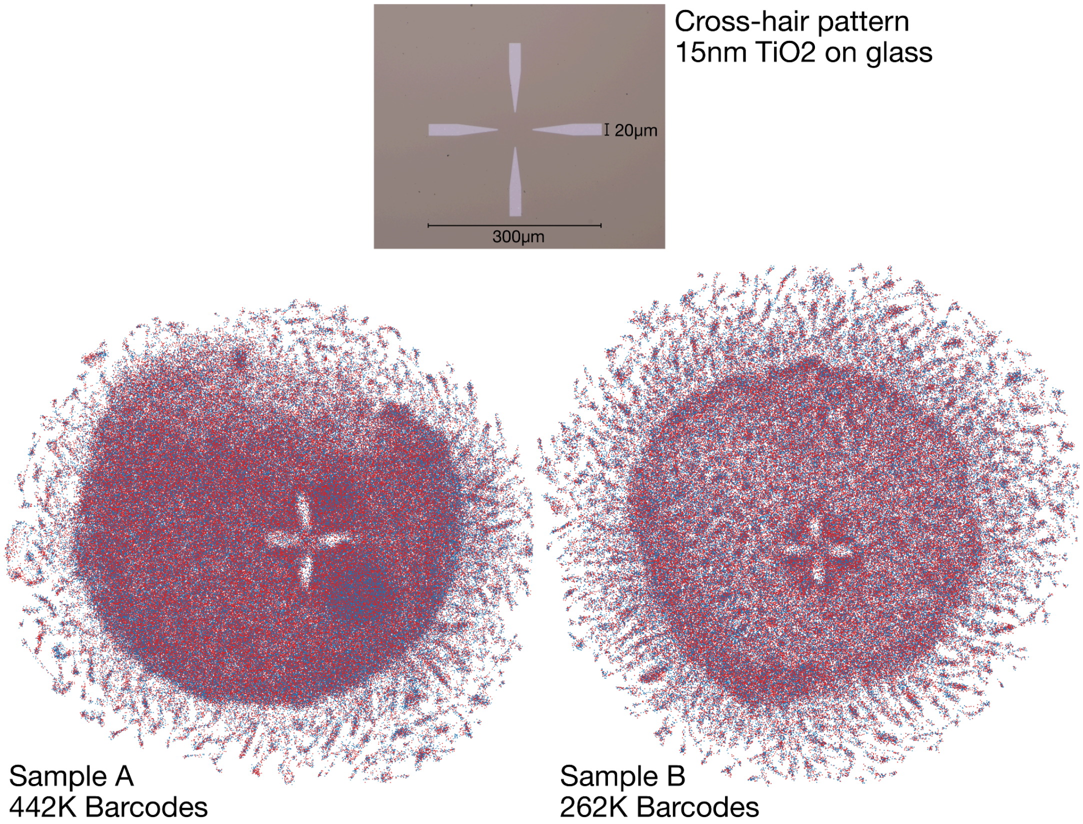


**Fig. S8: Additional examples of graph reconstruction from lithographically defined patterns.** Top: Optical micrograph of a crosshair pattern, 300µm in width with 20µm-wide arms, fabricated using the same microfabrication process described in the main text. Bottom: Graph reconstructions from two independent chips, both revealing gaps in the barcode distributions that reproduce the crosshair pattern.

| Label | Sequence |
|---|---|
| 3P | 5’-G*G*A*A*G*T*A*G*C*G*T*G*G*A*TGG WNNNWNNNWNNNNWNNWNNNWNNNN GGTGAGCAGTTGGTCGG TTTTTTTTTT-(DBCO)-3’ |
| 5P | 5’-(DBCO)-(TEG spacer)-TTTTTTTTT-(HEG spacer)-AGGTTAC AGAGTGCGGC WNNNWNNNNWNNNNWNNNNWNNNSW GGAAGTA GCGTGGATGG-(Inverted dT)-3’ |
| EG | 5’-(DBCO)-(TEG spacer)-TTTTTTTTTTTTTTT UTCUTCUTCUTCUTCUTCU CCGACTTCCTGACCAACTGCTCACC-3’ |
| PrimerFwd | AGGTTACAGAGTGCGGC |
| PrimerRev | CCGACTTCCTGACCAAC |

**Table S1: DNA sequences used in this study.** Asterisks (e.g. G*) represent phosphorothioated DNA bases, which act both as weak binding sites as well as nuclease protection.

| **Storage Technology** | **Volume ($nm^3$/bit)** |
|---|---|
| Sequence-based DNA storage (Theoretical minimum) | ~0.7 |
| Flash memory (3D NAND) | $\sim 1.1\times10^{6}$ |
| Magnetic tape (LTO-10) | $\sim 2.1\times10^{8}$ |
| Optical Archival Disc (500GB) | $\sim 3.3\times10^{9}$ |

**Table S2: Comparison of physical storage volume per bit across different storage technologies.** For conventional DNA storage, the molecular volume was calculated assuming 1.5 bits per base pair, a DNA diameter of 2nm, and an axial rise of 0.34nm per base pair. For SPAN, the volume was calculated assuming nine double-stranded DNA molecules of 100bp=(34nm) per stored bit. For 3D NAND flash, the volume was estimated using an areal density of 37 Gbit/$mm^2$ (*55*) and a minimum die thickness of 40µm. Magnetic tape volume was calculated using the physical dimensions and native capacity of Fujifilm's 40 TB LTO-10 tape (*56*). Optical storage volume was calculated using a Sony's 500GB Archival Disc (*57*) with a diameter of 120mm, 1.2mm thickness, and central hole diameter of 15mm.

**Movie S1:** Reconstruction of the MIT logo pattern as a function of sequencing reads
